# Early Deprivation Impairs Perforant Pathway Connectivity and Contextual Memory in Adolescent Male Mice

**DOI:** 10.1101/2023.08.08.552517

**Authors:** Rafiad Islam, Jordon D. White, Tanzil M. Arefin, Sameet Mehta, Xinran Liu, Baruh Polis, Lauryn Giuliano, Sahabuddin Ahmed, Christian Bowers, Jiangyang Zhang, Arie Kaffman

## Abstract

Early-life adversity causes reduced hippocampal volume and abnormal hippocampal connectivity and function, with evidence indicating more prominent deficits in males compared to females. Reelin-positive projections from the lateral entorhinal cortex (**LEC**) to the dorsal hippocampus are essential for encoding contextual and semantic memories in diverse mammalian species, including humans and rodents. However, the impact of early-life adversity on these projections and their contribution to hippocampal-dependent deficits have not been reported. Using a modified limited bedding (**LB**) mouse model of early adversity that extends the impoverished conditions from birth to postnatal day 25 (**P25**), we found severe impairment in contextual fear conditioning for adolescent LB male but not LB female mice. Using retrograde tracing, we found that the number of reeling-positive projections from the LEC to the dorsal hippocampus is significantly reduced in LB males but not LB females. Further, the number of projections was highly correlated with deficits in contextual memory and hypomyelination in perforant pathway terminals located in the dorsal hippocampus. Ex vivo high-resolution diffusion magnetic resonance imaging confirmed reduced structural connectivity between the entorhinal cortex and the dorsal hippocampus and revealed extensive cortical atrophy that resembled abnormalities reported in children exposed to severe deprivation. Given the essential role that reelin-positive projections play in contextual memory, these findings suggest a novel mechanism to explain the pronounced contextual memory deficits seen in LB males.

## INTRODUCTION

Early-life adversity (**ELA**) alters brain development and increases the risk for multiple psychiatric conditions (1, 2). Some of the most reproducible findings in individuals exposed to ELA are reduced hippocampal volume and abnormal hippocampal connectivity and function, which are often more prominent in males compared to females (1, 3-6). The mechanisms responsible for these changes are yet to be clarified, but are likely due to sex-differentiated structural and functional changes early in life (7, 8) that are sensitive to key components of ELA such as stress and levels of sensory, social, and cognitive stimulation (9-12).

Work in rodents can clarify essential details regarding the underlying biology by which ELA alters hippocampus development and function in male and females. This assertion is supported by work showing that hippocampal development is conserved across mammalian species (13-15) and that early adversity impairs hippocampal development and long-term function in rodents (16). Further, elegant work in rodents has identified several mechanisms by which ELA in the form of limited bedding and nesting impairs hippocampal developmental and function (11, 12, 17). Despite substantial progress, key questions remain. For example, most studies to date have focused on outcomes in adult males, with only a handful of examples examining this issue in adolescent male and female rodents (12, 16, 17). Second, the commonly used limited bedding and nesting paradigm takes place from postnatal day 2 (**P2**) to P9 (18). This period is equivalent to developmental processes in the human hippocampus that extends from the third trimester to roughly 1-3 years of age, precluding key developmental processes that take place during childhood and adolescence including dentate gyrus maturation, synaptogenesis, dendritic arborization, synaptic pruning, GABAergic input, and myelination (19-22). Finally, little attention has been paid to the possibility that abnormal connectivity involving key cortical regions, such as the entorhinal cortex, may contribute to aberrant hippocampal development and function in individuals exposed to ELA. These connections originate from reelin-positive cells in layer 2 of the lateral entorhinal cortex (**LEC**) and the medial entorhinal cortex (**MEC**) to form the perforant pathway, the primary input into granule cells in the dentate gyrus (**DG**) and to a lesser extent pyramidal neurons in CA2 and CA3 subregions of the hippocampus (23). Mature perforant pathway terminals are visible relatively early in development, around postnatal day 10 (P10) and P14 in mice and rhesus monkeys, respectively (24-26). Functional connectivity between these terminals and granule cells in the DG undergo notable changes during childhood and adolescence, including myelination and expansion of synaptic input onto granule cells (22, 26). LEC input targets newly formed neuroblasts, thereby serving an important role in guiding postnatal neurogenesis and pattern separation (27). Disruption of perforant connectivity impairs episodic memory in juvenile rodents (28) and children (29), underscoring an essential role for these connections in normal hippocampal development and function.

To address these issues, we have extended the limited bedding LB procedure from P0 (birth) to P25 and tested the impact of this manipulation on hippocampal function and connectivity in adolescent male and female mice. We show that LB causes more severe deficits in contextual memory in adolescent male mice compared to females and that this effect is at least partly due to sex-specific reduced connectivity between the LEC and the DG. A failure of reelin-positive cells to project from the LEC to the dorsal hippocampus is responsible for the reduced perforant pathway connectivity in males and contributes to deficits in myelination of terminals located in the stratum lacunosum moleculare (**SLM**). Finally, both adolescent LB male and female mice show hyperactivity and extensive reduction in both cortical and subcortical grey matter that resemble abnormalities reported in children and adolescents exposed to severe deprivation (30-34).

## METHODS

### Animals

BALB/cByj mice (Stock # 001026) were kept on a standard 12:12 hr light-dark cycle (lights on at 7:00AM), with food provided *ad libitum*. Temperature and humidity were held constant (23 ± 1°C and 43% ± 2) and background noise in the room was kept at dB: 56.5. All studies were approved by the Institutional Animal Care and Use Committee (IACUC) at Yale University and were conducted in accordance with the recommendations of the NIH Guide for the Care and the Use of Laboratory Animals.

### Limited Bedding (LB)

The limited bedding (LB) procedure was done as described previously (10, 35). Briefly, breeding cages were set up using a 3:1 female to male harem in standard mouse Plexiglas cages with 2 cups corncob bedding and no nesting material. Visibly pregnant dams were transferred to ‘maternity cages’ containing 2 cups corncob bedding with no nesting material and 3 chow pellets on the floor. At birth (P0), litters were culled to 5–8 pups and randomized to either control (CTL) or limited bedding (LB). CTL litters were provided with 500 cc of corncob bedding, 15cc of soiled bedding from the birth cage, and one 5 × 5 cm nestlet from P0-25. LB litters were provided with 125 cc corncob, 15 cc of soiled bedding from the birth cage, and no nestlet from P0-25. Bedding was changed on P7, P14, and P21. Mice were weaned on P26 and housed with 2-3 same-sex littermates per cage with 500cc of corncob bedding, no nesting material, and 2-3 chow pellets on the floor.

### Tissue Collection and Processing

Tissue collection for immunohistochemistry was done as described previously (10, 35). To characterize RNA, DNA, and protein content in the hippocampus, P17 pups were anesthetized, and the right and left hippocampi were dissected, snap-frozen in liquid nitrogen, and stored at - 80 °C until further processing. The left hippocampus was homogenized on ice in the lysis buffer from the AllPrep^®^ DNA/RNA/Protein Mini Kit (QIAGEN, Hilden, Germany). A sample from the lysate was used to quantify total DNA and RNA content using the appropriate Qubit^®^ Fluorometer kit (Cat. # Q32850 for DNA and Q32852 for RNA). The remaining homogenate was then processed according to manufacturer’s instructions to purify RNA, DNA, and proteins. Samples were stored at -80°C until further processing. Purified total protein was quantified using Pierce™ BCA Protein Assay Kit (Cat. # 23227).

### RNA-seq

RNA samples from the left hippocampus of P17 pups (CTL males = 6, CTL females = 9, LB males = 7, LB females = 9 from 4-6 independent litters) were sequenced at the Yale Center for Genomic Analysis. RNA quality and concentration were assessed using the 2100 Bioanalyzer on an RNA 6000 Nano Assay (Agilent Technologies, Inc, all RINs > 9). Libraries were constructed by amplifying 500 ng of total RNA using 9 PCR cycles with the KAPA mRNA HyperPrep Kit (KAPA Biosystems). cDNA libraries were validated using the Bioanalyzer 2100 on a High Sensitivity DNA assay and quantified using the KAPA Library Quantification Kit for Illumina Platforms on a Roche Lightcycler 480. Sequencing was done on an Illumina NovaSeq 6000 using the S4 XP workflow 2x100 with each library using 1.25% of the S4 lane. Reads were trimmed for quality using custom scripts with a minimum accepted length of 45 bases. The trimmed reads were then aligned to the mm10 reference mouse genome using gencode annotation (36) and HISAT2 software (37). Transcripts per million (TPM) estimation were calculated using StringTie (37).

### Immunohistochemistry

Perfused brains were sectioned coronally using a VT1000S vibratome (Leica) to obtain 50-micron sections and sorted into 6 pools, each containing 16–18 slices spaced at 240-micron intervals that systematically span the entire rostral-caudal axis of the brain. Oligodendrocyte differentiation was assessed by incubating free-floating sections overnight at room temperature with rabbit PDGFRα (D1E1E) XP^®^ antibodies (Cell Signaling, cat #3174,1:200), mouse anti-CC1 (BD Biosciences, cat #558774, 1:100), and rat anti-MBP (BIO-RAD, cat # MCA409S, 1:500). Axonal staining and myelination in the SLM were assessed with rat anti MBP (BIO-RAD, cat # MCA409S 1:500) and mouse anti-Neurofilament H (NF-H), Phosphorylated Antibody (BioLegend, Cat# 8016021, 1:1000). Number of reelin positive cells was assessed using a mouse anti-reelin antibody (Cat # AB78540-1001, 1:400). Slices were then incubated with the appropriate fluorescently labeled secondary antibodies (ThermoFisher, Cat #A-21422, # A-11008, # A-21094, 1:200) and mounted on glass slides with VECTASHIELD HardSet antifade mounting medium with DAPI (Vector laboratories Cat# 10955).

### Stereology, Cell Counting, and Image Analysis

Stereological analysis was done by tracing the borders of the granule cell layer of the dentate gyrus (DG) and the pyramidal cell layer of the Cornu Ammonis (**CA**) regions of the hippocampus under low magnification (-2.5x) using the Stereo Investigator 10 software (MBF Bioscience) and volume-was calculated using the Cavalieri’s principle (38, 39). To quantify PDGFRα and CC1 cell number and MBP intensity, a 450 × 250 µm square area containing the SLM was selected and analyzed using Imaris 10.0 (Bitplane, Oxford Instruments). The object selection spots model was used to analyze PDGFRα and CC1 with cell diameter set to 6.5 µm for PDGFRα and 13.5 µm for CC1 and Z diameter kept as default. MBP and NF-H intensities were determined using the object selection surface model using 0.720 µm threshold with the intensity max filter applied. Colocalization between MBP and NF-H was done with the Object-Based-Colocalization model using the shortest distance to surfaces of 0.1 µm. Three slices were counted per animal and averaged to determine the number of PDGFRα- and CC1-positive cells, MBP intensity, and MBP/NF-H colocalization for each mouse. For counting CTB-positive cells, a 200 X 300 μm square area containing layers 2-3 of the LEC was cropped and the number of cells counted in Imaris using the Spots Model with XY diameter of 20 μm, default Z-diameter, and max intensity filter settings. 5 slices were counted per animal and averaged to obtain the number of reelin-positive cells in the LEC for each animal.

### Electron Microscopy

P17 mice were perfused with cold PBS followed by 4% paraformaldehyde in PBS. Brains were then dissected out and immersed in a solution containing 2.5% glutaraldehyde and 2% paraformaldehyde in 0.1M sodium cacodylate buffer (pH 7.4) and kept overnight at 4 °C. 200 micron-thick sections containing the dorsal hippocampus (Paxinos coordinates, Bregma= - 1.34mm to -2.80mm) were cut using a VT1000S vibratome (Leica) and post-fixed with 1% OsO_4_, 0.8% potassium ferricyanide at room temperature for one hr. Specimens were then *en bloc* stained with 2% aqueous uranyl acetate for 30 min, dehydrated in a graded series of ethanol to 100%, substituted with propylene oxide, and embedded in EMbed 812 resin. Sample blocks were polymerized in an oven at 60 °C overnight. To locate SLM, semithin sections (250 nm) were prepared and stained by the solution of toluidine blue, 1% sodium borate. TEM thin sections (60 nm) were obtained using a Leica ultramicrotome (UC7) and post-stained with 2% uranyl acetate and lead citrate. Sections were examined with a FEI Tecnai transmission electron microscope at 80 kV accelerating voltage, and digital images recorded with an Olympus Morada CCD camera and iTEM imaging software. The MyelTracer software (40) was used to determine axonal diameter and G-ratio defined as the inner axonal diameter divided by the diameter of the axon plus the myelin sheath that encases it, n = 20 axons per mouse and n = 3 mice per group.

### Behavior

Open field - (35, 41) and contextual fear conditioning (41) were done as previously described except that acoustic cue conditioning was not tested on the third day of the contextual fear conditioning. Twenty-four hours after behavioral testing, mice were perfused for immunohistochemistry or diffusion magnetic resonance imaging (**dMRI**).

### Ex vivo dMRI

P29 Adolescent CTL and LB mice (N = 6 mice per rearing and sex, a total of 24 mice) were perfused and processed for high-resolution ex vivo dMRI using a 7-Tesla MR system equipped with a 4-channel cryogenic probe as described previously (41, 42). Whole-brain-voxel-based morphometric analysis was used to identify local volumetric changes affected by rearing, sex, and their interaction (2 x 2 ANOVA, FDR corrected, α = 0.1, p < 0.0105, cluster size > 25 voxels). Fiber tractography was used to assess structural connectivity between the entorhinal cortex and the dorsal hippocampus following previously described methodology (41-43).

### Retrograde Labeling

P29 mice were anesthetized with isoflurane (1–3%) and 50 nl of the retrograde tracer cholera toxin subunit B (5% wt./vol in PBS, ThermoFisher Scientific, Cat # C-34775) were injected at 10nl/sec into the left SLM (ML: -1.2, AP: -2, DV: -1.9) using a 5 uL Hamilton syringe equipped with a 32 blunt needle (Hamilton, Cat# 7803-04, 0.5 inch, PST 3). After injection, the needle was left in place for 5 min and then slowly removed. Incision was closed with Vet Bond and mice were allowed to fully recover prior to being returned to their home cage.

### Statistical Analysis

Statistical analyses were done using Matlab R2022b (www.mathworks.com), SPSS (Version 26.0. Armonk, NY: IBM Corp.), and visualized with GraphPad Prism 9.0 - (GraphPad Software, La Jolla California USA). Data were examined using a 2 X 2 ANOVA with rearing condition (CTL, LB) and sex as fixed factors followed by Tukey’s HSD post-hoc test for significant rearing by sex interaction. RNA-seq analysis was conducted by first removing genes with very low expression (average TPM < 3 in all three groups). This was done to increase statistical power and focus on genes that are more likely to drive developmental changes. Counts of the 13,245 genes that passed the initial screen were analyzed using DESeq2 script in R (44) to identify genes that are differentially expressed between CTL and LB using Benjamini-Hochberg false discovery rate (FDR) < 0.05. All raw data and analyses are available at GEO archive accession # <u>GSE142305</u>. Enrichment and transcription factor analyses were conducted using MetaCore™ (Clarivate Analytics) workflows on gene lists that passed a corrected p-value of < 0.05.

## RESULTS

### LB Inhibits Oligodendrocyte Progenitor-Cell Differentiation in the SLM Region of the Developing Hippocampus

We have previously shown that exposure to LB reduces hippocampal size and protein content in P14 pups (35). We replicated and extended these findings in P17 pups by showing that LB reduces the size of the granule cell layer (**GCL**), Cornu Ammonis (**CA)** pyramidal layer, DNA, RNA, and protein content in male and female pups (**Fig S1**). Bulk RNA-seq was used to further elucidate the mechanisms by which LB alters hippocampal development (**Fig 1A**) and identified 797 differentially regulated genes (**DRG**) between CTL and LB (FDR < 0.05), most of which had a small effect size (i.e., of log2 < 0.4, green/turquoise heat map, **Fig 1B**). Pathway analysis identified oligodendrocyte differentiation and myelination as the most significantly downregulated pathways in LB, whereas cytoskeleton remodeling was the most upregulated pathway in LB (**Fig 1C**). The 20 most up- and down-regulated genes are shown in **Fig S2**.

**Figure 1.**
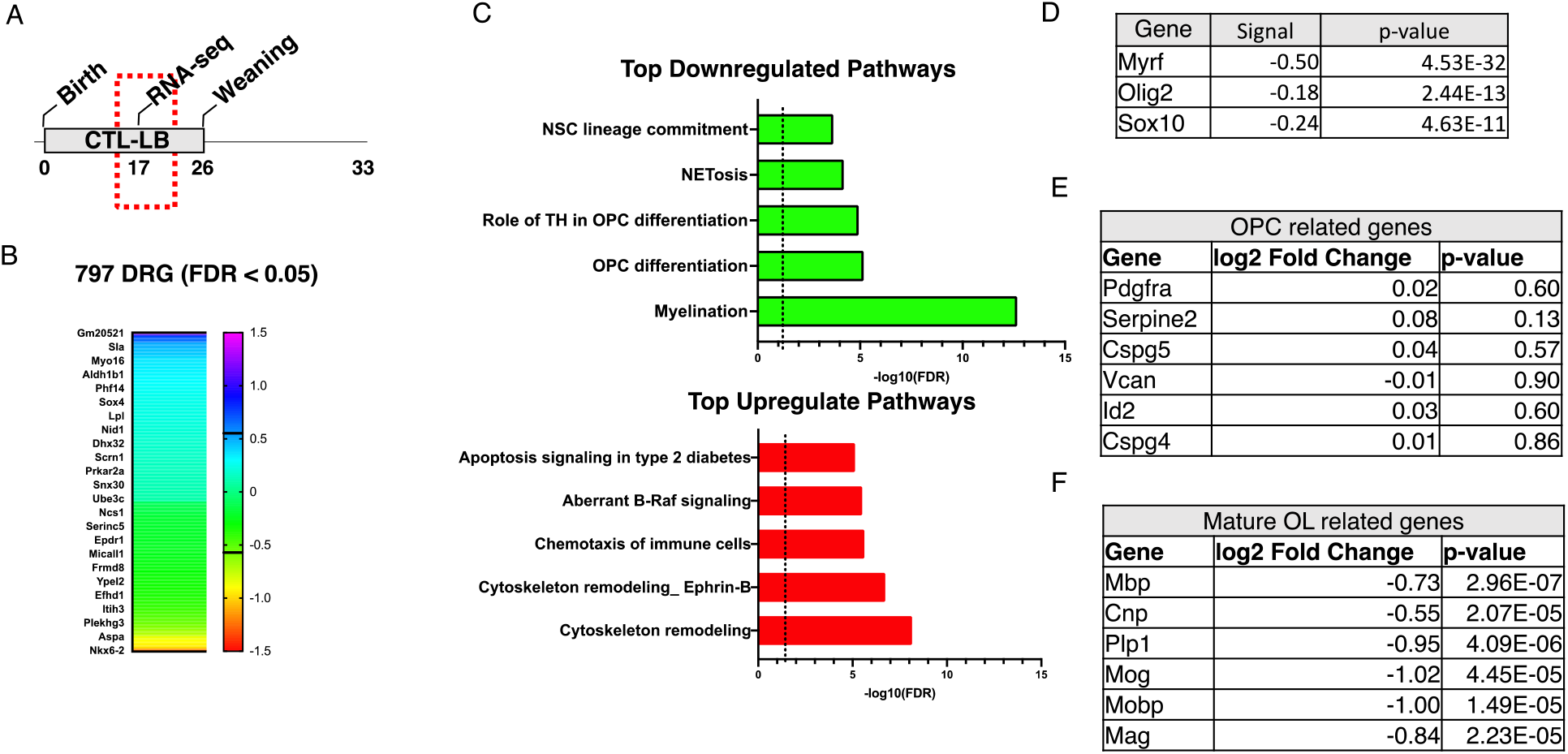
LB Inhibits Oligodendrocyte Differentiation in the Hippocampus of P17 Pups. (A) Experimental timeline. (B) Heat map showing effect size of differentially regulated genes (DRG) at FDR < 0.05 between CTL and LB. (C) Pathway analysis of top down-regulated and up-regulated pathways affected by LB. (D) Transcription factor analysis identified changes in Myrf, Olig2 and Sox10 activity as the most dysregulated factors affected by LB. LB does not affect expression of OPC-specific genes (E) but significantly decreases expression of genes expressed in mature oligodendrocytes (F).

Transcription factor analysis identified reduced activity of Myrf, Olig2, and Sox10 as the key regulators of abnormal myelination in LB mice (**Fig 1D**). Since Myrf is necessary for oligodendrocyte differentiation (45, 46) we assessed the effects of LB on genes that are specifically expressed in oligodendrocyte progenitor cells (**OPCs**) and mature oligodendrocytes (**OL**). LB had no impact on OPC-specific genes (**Fig 1E**), but significantly reduced expression of OL-specific genes (**Fig 1F**), indicating that LB impairs the differentiation of OPC in the developing hippocampus.

To further validate the RNA-seq data, we assessed the effects of LB on oligodendrocyte maturation in the stratum lacunosum moleculare (**SLM**), one of the most myelinated regions in the developing hippocampus (**Fig 2A**) (22, 47, 48). This was done by quantifying levels of MBP as a marker of myelination, number of PDGFRα-positive cells (OPCs), and number of CC1 positive cells (OL) in the SLM of P17 pups. Consistent with the RNA-seq data, we found a significant reduction in MBP staining in LB compared to control (F (1, 16) = 15.42, P = 0.0012, ηp^2^ = 0.49) (**Fig 2B-C**). There was also a significant reduction in the number of PDGFRα-positive cells in LB pups (rearing effect: F (1, 16) = 5.88, P = 0.0275, ηp^2^ = 0.27) (**Fig 2 B & D**). For CC1-postive cells, there was a significant effect of rearing (F (1, 16) = 35.56, P<0.0001, ηp^2^ = 0.69) and sex (F (1, 16) = 4.664 P = 0.046) but no significant interaction between rearing and sex (**Fig 2E**). Since LB reduced both OPC and OL in the SLM, an OPC differentiation index was generated by dividing the number of CC1-positive cells by the number of PDGFRα positive cells. There was a significant reduction in the OPC differentiation index in LB mice (F (1, 16) = 32.4, P< 0.000, ηp^2^ = 0.67, **Fig 2F**) indicating that impaired OPC differentiation is primarily responsible for the reduced myelination in the SLM.

**Figure 2.**
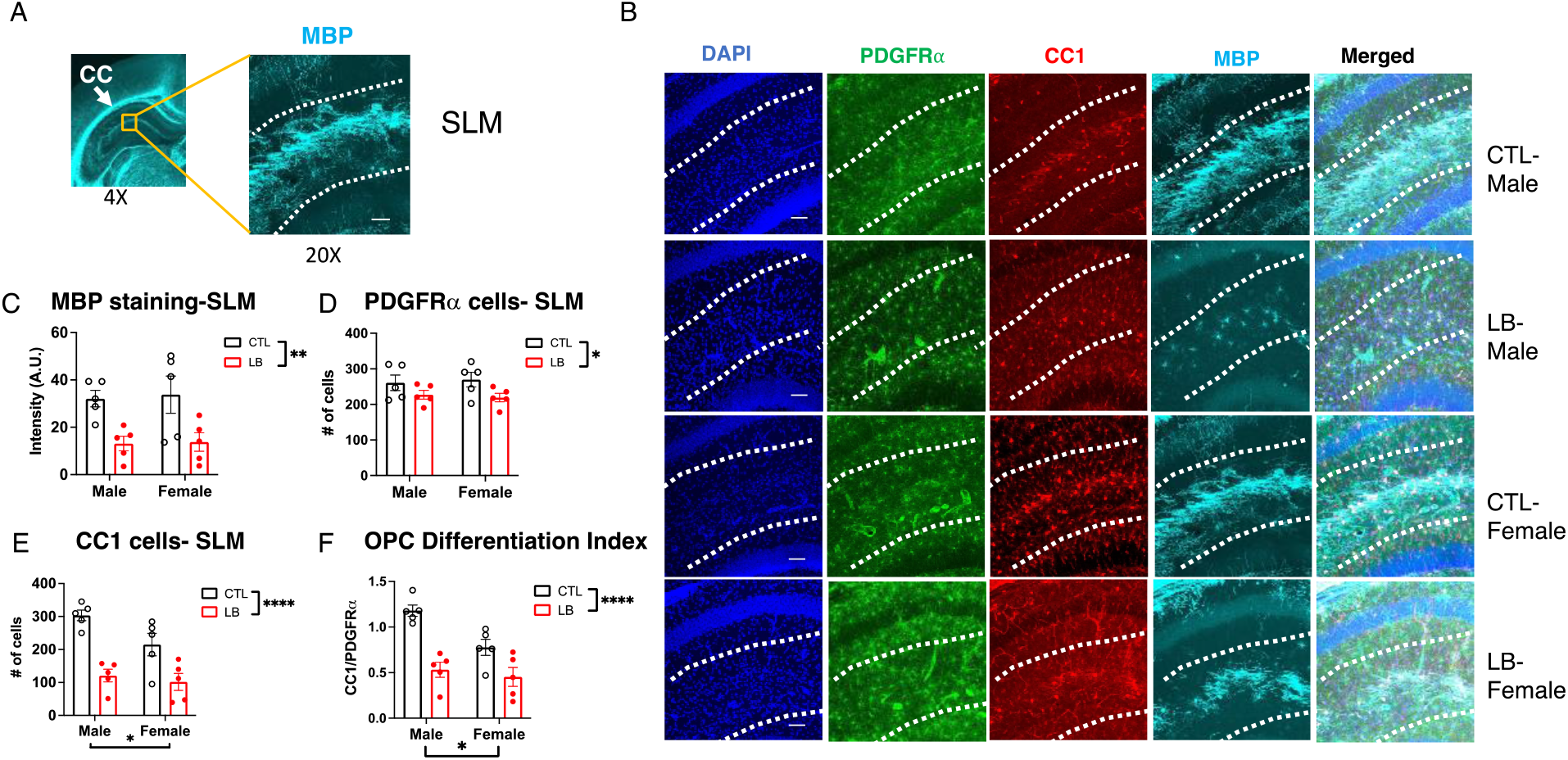
OPC Differentiation in the SLM is Impaired in P17 LB mice. (A) Low (4X) and higher (20X) magnification of myelin basic protein (MBP) staining in the SLM of P17 pups. (B) Representative images of PDGFRα-positive OPC, CC1-positive mature oligodendrocytes, and MBP in the SLM. Quantification of MBP (C), PDGFRα-positive OPCs (D), CC1 mature oligodendrocytes (E) and OPC differentiation index (F) in the SLM. N= 5-6 mice per rearing and sex group. Error bars represent mean ± SEM. *p< 0.05, **p< 0.01, ****p< 0.0001.

Using electron microscopy, we further confirmed reduced axonal myelination in the SLM of 17-day old LB pups (**Fig 3A**). LB mice showed significant increase in the G-ratio (t (118) = 15.57, P < 0.0001, Cohen’s d = 2.84, **Fig 3B**). Axonal diameter was also increased in LB compared to CTL animals (CTL = 1.05 ± 0.04 um, LB = 1.47 ± 0.074 um, t (95) = 4.87, P< 0.001, Cohen’s d = 0.89), indicating that the reduced myelination was not due to reduced axonal diameter. Further, the increase in G-ratio seen in LB continued to be significant even after adjusting for the increase in axonal diameter (ANCOVA, F=130.759, P < 0.001, **Fig 3C**), demonstrating that the elevated G-ratio seen in LB mice is primarily driven by reduced myelination.

**Figure 3.**
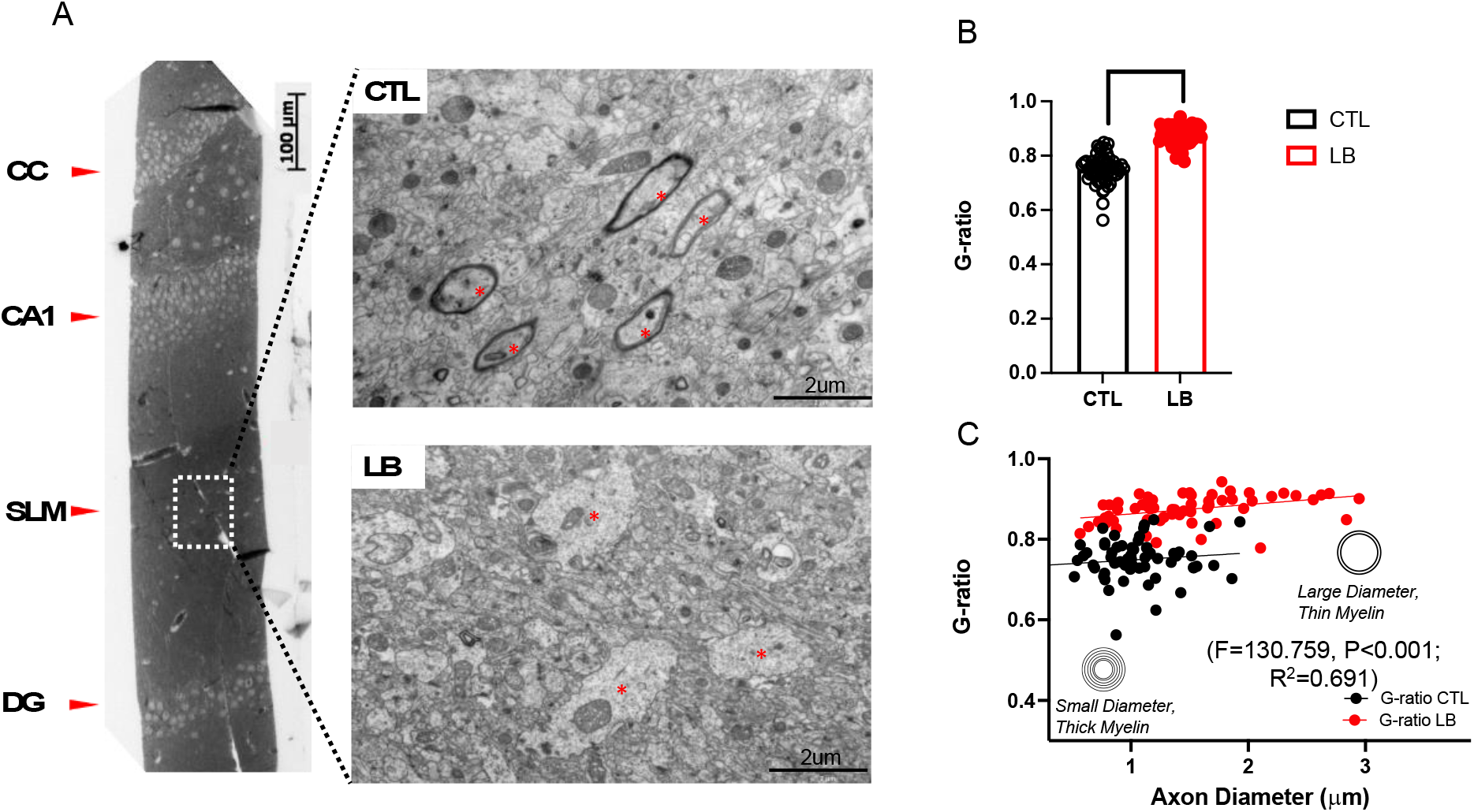
Transmission electron microscopy of myelinated axons in the SLM of P17 CTL and LB mice. (A) Left, a light microscopy image of toluidine-stained same sample to identify the SLM; Right, TEM images of the SLM in CTL and LB mice (visible axons indicated with red dots). (B) Image analysis of co-variance between G-ratio and axonal diameter (C). N = 20 axons per mouse with 3 male mice per condition. Error bars represent mean ± SEM. ****p < 0.0001.

### Axonal Staining is Reduced in the SLM of P17 LB mice

To test whether the reduced myelination is due to abnormal axonal innervation, we probed the hippocampus with anti-phosphorylated neurofilament H antibodies (NF-H) to quantify axonal fibers in the SLM of P17 control and LB mice (**Fig 4A**). There was a significant reduction in NF-H staining in LB mice (F (1, 16) = 11.38, P=0.0039), with no significant effect of sex and a trend for rearing by sex interaction (F (1, 16) = 3.30, P = 0.088) due to a greater reduction in NF-H staining in LB males compared to LB females (**Fig 4B**). A similar pattern was seen for MBP staining, with reduced myelination in LB males (P < 0.0001) but not in females (P= 0.48, interaction: F (1, 16) = 14.20, P = 0.0017, **Fig 4C**). The percent of neurofilaments co-stained with MBP was reduced in LB compared to CTL -(F (1, 16) = 5.78, P= 0.029) with no significant effects of sex or interaction (**Fig 4D**). These results suggest that other mechanisms, in addition to reduced axonal availability, contribute to the abnormal myelination seen in 17-day old LB mice. This assertion is consistent with the presence of large hypomyelinated axons in the SLM at this age (**Fig 3A**).

**Figure 4.**
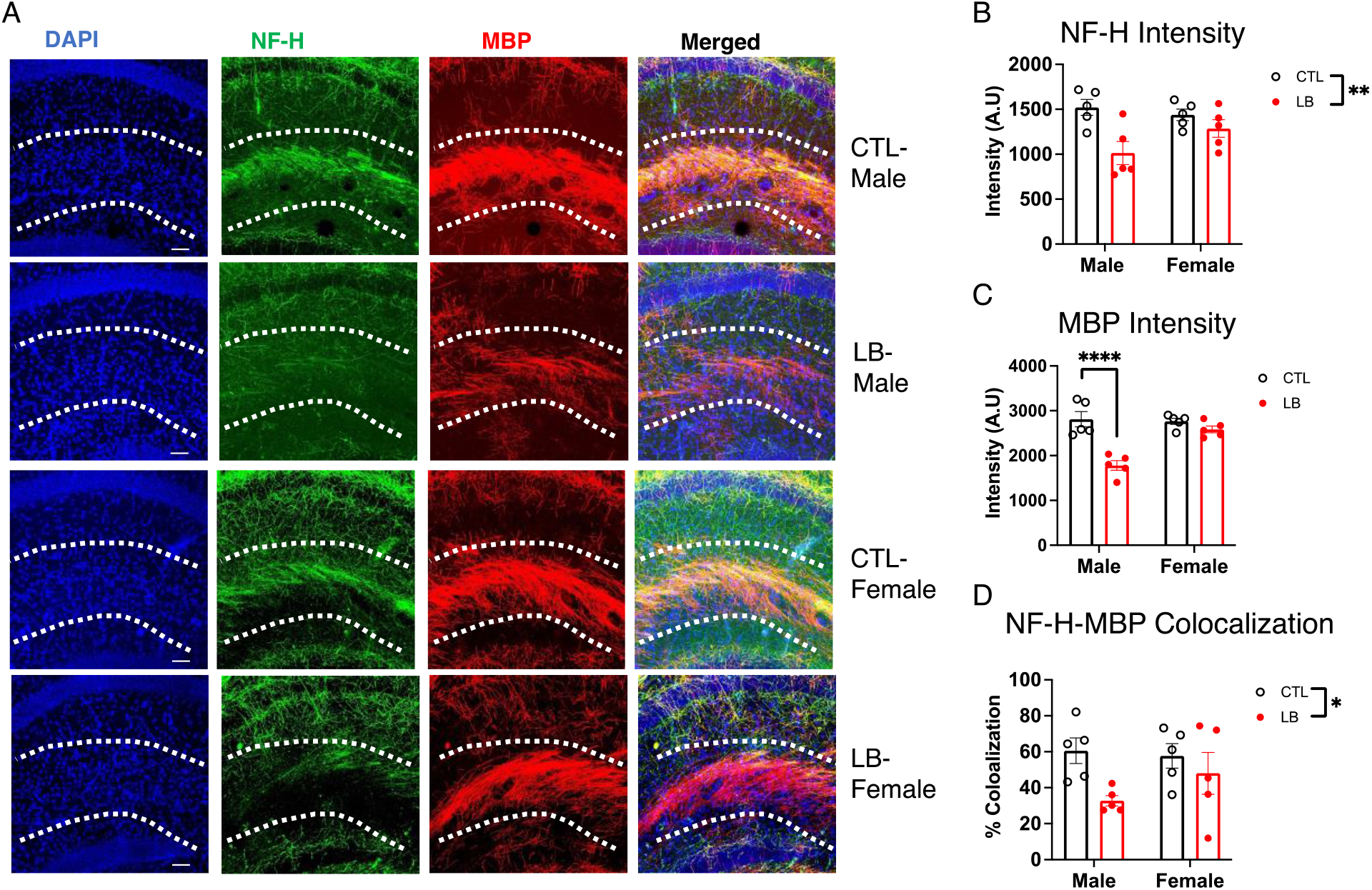
LB Causes a More Pronounced Reduction in Axonal Staining in the SLM of 17-day old LB Male Mice. (A) Representative images of axonal staining using anti-NF-H and anti-MBP antibodies in the SLM. Quantification of NF-H staining (B), MBP staining (C) and NF-H-MBP colocalization (D). Scale bars in Fig 4A are 50 microns. N = 5 mice per rearing and sex. Error bars represent mean ± SEM. *p < 0.05, **p < 0.01, ****p < 0.0001.

### Reduced Axonal Innervation Persists in P33 Adolescent Mice and Correlates with Deficits in Contextual Fear Conditioning

Next, we tested whether the reduction in myelination and axonal staining seen in the SLM of P17 pups persisted and contributed to hippocampal-dependent deficits in adolescent mice. This was done by first testing contextual freezing in P31-33 mice and then assessing axonal staining and myelination in the SLM (**Fig 5A**). Mice showed the expected increase in freezing in response to shock on the first day of training (F (2.958, 136.1) = 208, P< 0.0001). However, there was a significant interaction between rearing and sex for contextual freezing during the second day (F (1, 62) = 3.994, P = 0.05), driven by a significant reduction in freezing behavior in LB males (P= 0.0008) but not LB females (P= 0.43, **Fig 5B**).

**Figure 5.**
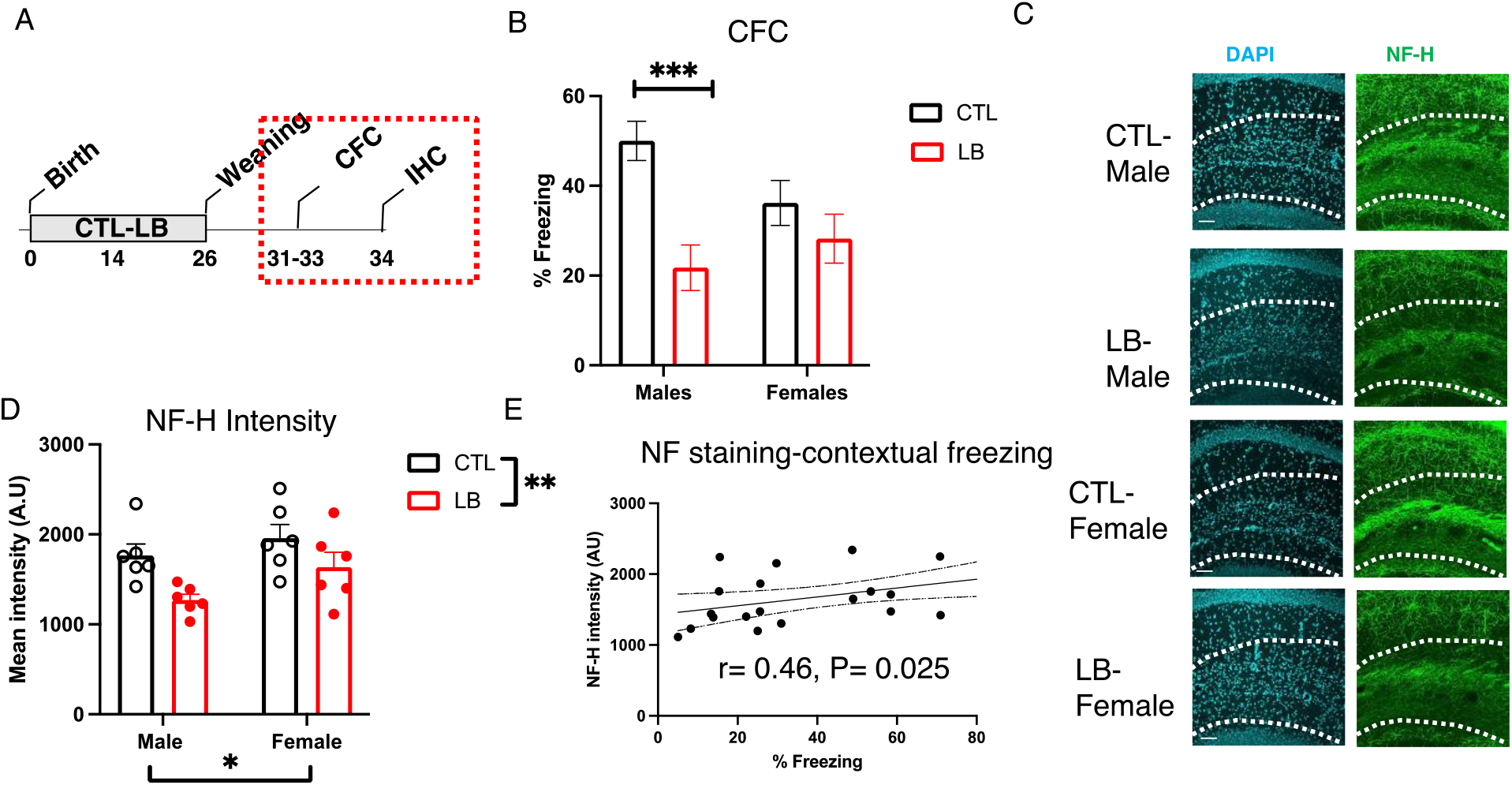
Reduced Axonal Staining in the SLM Persists in Adolescent LB Mice and Correlates with Contextual Freezing Behavior. (A) Experimental timeline. (B) Contextual fear conditioning CTL male = 17, CTL females = 18, LB males = 13, LB females = 18. (C-D) NF-H axonal staining in the SLM of adolescent mice, N = 6 mice per rearing and sex group. (E) Intensity of axonal staining correlates with contextual freezing behavior. Error bars represent mean ± SEM. *p< 0.05, **p< 0.01, ***p< 0.001.

Examination of NF-H staining in the SLM revealed significant effects of rearing (F (1, 20) = 9.55, P = 0.0058 and sex (F (1, 20) = 4.46, P = 0.047), but no significant interaction F (1, 20) = 0.426, P = 0.52 (**Fig 5C**). Nevertheless, as with the axonal staining at P17 (**Fig 4**), the reduction in axonal staining was more pronounced in LB males (Cohen’s d = 2.01) compared to LB females (Cohen’s d = 1.27) and the intensity of staining was significantly correlated with contextual freezing (**Fig 5E**). No significant effects of rearing, sex, or interaction were seen for the number of PDGFR1α progenitor cells or CC1 mature oligodendrocytes in the SLM (**Fig S3 A-C)**. However, there was a significant reduction in MBP staining in the SLM of LB adolescent mice (F (1, 19) = 4.40, P= 0.049) that was again more pronounced in males compared to females (**Fig S3D**). Together, these results indicate that reduced axonal staining and myelination persist in the SLM of adolescent LB mice and may contribute to the sex-specific deficits seen in contextual fear conditioning -

### Adolescent LB Mice Are Hyperactive and Show Extensive Cortical Atrophy

We have recently proposed that LB is a mouse model of childhood deprivation and neglect (6, 10). Given that hyperactivity and extensive cortical thinning are consistent findings in children and adolescents exposed to severe social and cognitive deprivation (49, 50) we used the open field test and high-resolution dMRI to assess these two outcomes in P29 mice. LB male and female mice showed a significant increase in locomotor activity in the open field test (F (1, 65) = 3.95 P = 0.05) with no significant effect of sex or interaction (**Fig 6A)**. There was a significant interaction between rearing and sex for the time spent in the center (interaction: F (1, 65) = 7.71, P = 0.0072, **Fig 6B**) that was due to a non-significant reduction in time spent in the center in LB males (P= 0.19) and a trend for an increase time in the center for LB females compared to CTL (P = 0.051). These findings replicate our previous work in adolescent LB mice (35) and confirm increased hyperactivity in LB adolescent male and female mice. Next, we conducted a 2 X 2 whole-brain voxel-based analysis to identify volumetric changes induced by rearing, sex, and their interaction (41, 42). This unbiased approach revealed extensive volumetric reduction in LB adolescent male and female mice that included sensory and motor cortex, prefrontal cortex, hippocampus, amygdala, and thalamus (shown in red in **Fig 6C**). Follow-up region of interest analysis confirmed reduced CA1 (F (1, 20) = 15.25, P = 0.0009, **Fig 6D**) and dentate gyrus volumes (F (1, 20) = 13.37, P = 0.0016, **Fig 6E**) in both LB male and female mice. These findings indicate that the lower hippocampal volume, seen in P17 LB pups (**Fig S1**), persists in adolescent mice and is likely to represent a global atrophy across multiple cortical and subcortical grey matter regions. Few brain regions showed a significant effect of sex (**Fig S4**), and the only brain region that showed significant rearing by sex interaction was the left entorhinal cortex **(Fig 6F**). A region of interest analysis confirmed significant sex-by-rearing interaction in the left entorhinal cortex (F (1, 20) = 4.397, P = 0.049) driven by reduced volume in LB males (P = 0.0015), but not LB females (P = 0.54, **Fig 6G**). A similar analysis in the right entorhinal cortex found significant rearing effect (F (1, 20) = 11.85, P = 0.0026), but no significant impact of sex (F (1, 20) = 0.61, P= 0.44) or interaction (F (1, 20) = 1.39, P= 0.25, **Fig 6H**).

**Figure 6.**
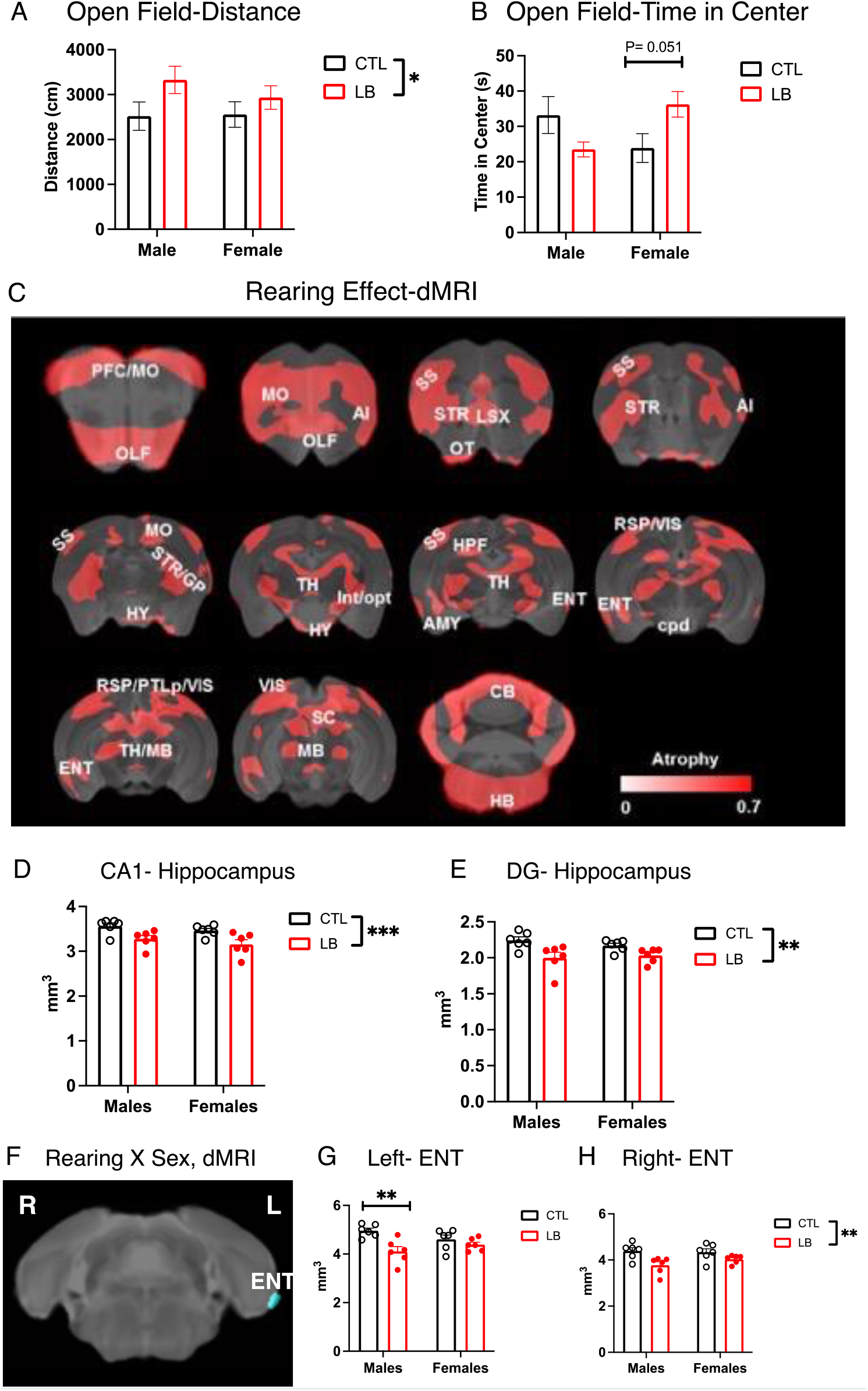
Adolescent LB Mice Are Hyperactive and Show Extensive Reduction in Cortical and Subcortical Grey Matter. Distance traveled (A) and time in the center (B) in the open field test in P29 mice (CTL males N = 15, CTL females N = 14, LB males N = 16, LB females N = 24). (C) Effect of rearing on volumetric changes (Minimal cluster size > 25 voxels, FDR < 0.1, p < 0.005). Areas with significant volume reduction are shown in red (N = 6 per rearing and sex). Effect of LB on CA1 (D) and dentate gyrus (E) volumes. (F-H) The left entorhinal cortex was the only region showing significant rearing by sex interaction (Minimal cluster size > 5 voxels, FDR < 0.1, p < 0.0023). Abbreviations: AI: Insular cortex, CB: Cerebellum, cpd: cerebal peduncle, ENT: Entorhinal cortex, GP: Globus pallidus, HB: Hindbrain, HPF: Hippocampus, HY: HY: Hypothalamus, Int: Internal capsule, LSX: Lateral septal complex, MO: Motor cortex, OLF: Olfactory area, opt: Optic tract, OT: Olfactory tubercle, PFC: Prefrontal cortex, PTLp: Posterior parietal association cortex, RSP: Retrosplenial cortex,, SC: Superior colliculus, SS: Somatosensory cortex, STR: Striatum, TH Thalamus, VIS: Visual cortex.

### LB Reduces Reelin-positive Projections in Male Mice

Most axonal terminals in the SML originate from the entorhinal cortex (22) suggesting that reduced axonal staining in the SML (**Fig 5D**) reflects abnormal connectivity between these two brain regions in LB adolescent mice. Indeed, dMRI tractography revealed a significant reduction in structural connectivity between the entorhinal cortex and the dorsal hippocampus on the left (F (1, 20) = 8.241, P = 0.0095, **Fig 7A-B**) and the right hemispheres (F (1, 20) = 5.659, P = 0.0274, **Fig 7C**). Although there were no significant effects of sex or interaction between rearing and sex, the effect sizes in males were larger compared to females (Cohen’s d left: M = 1.39, F = 0.92; Cohen’s d right: M = 1.65, F = 0.39). To further validated these findings, we administered a fluorescently labeled retrograde tracer Alexa-555-CTB into the left SLM of P28 CTL and LB mice, tested contextual fear conditioning at P34, and processed the mice a day later to assess the number of CTB-positive cells in the left entorhinal cortex (**Fig 8A**). LB mice showed a significant reduction in contextual freezing (F (1, 23) = 6.80, P = 0.016) that was more pronounced in males compared to females (**Fig 8B**).

**Figure 7.**
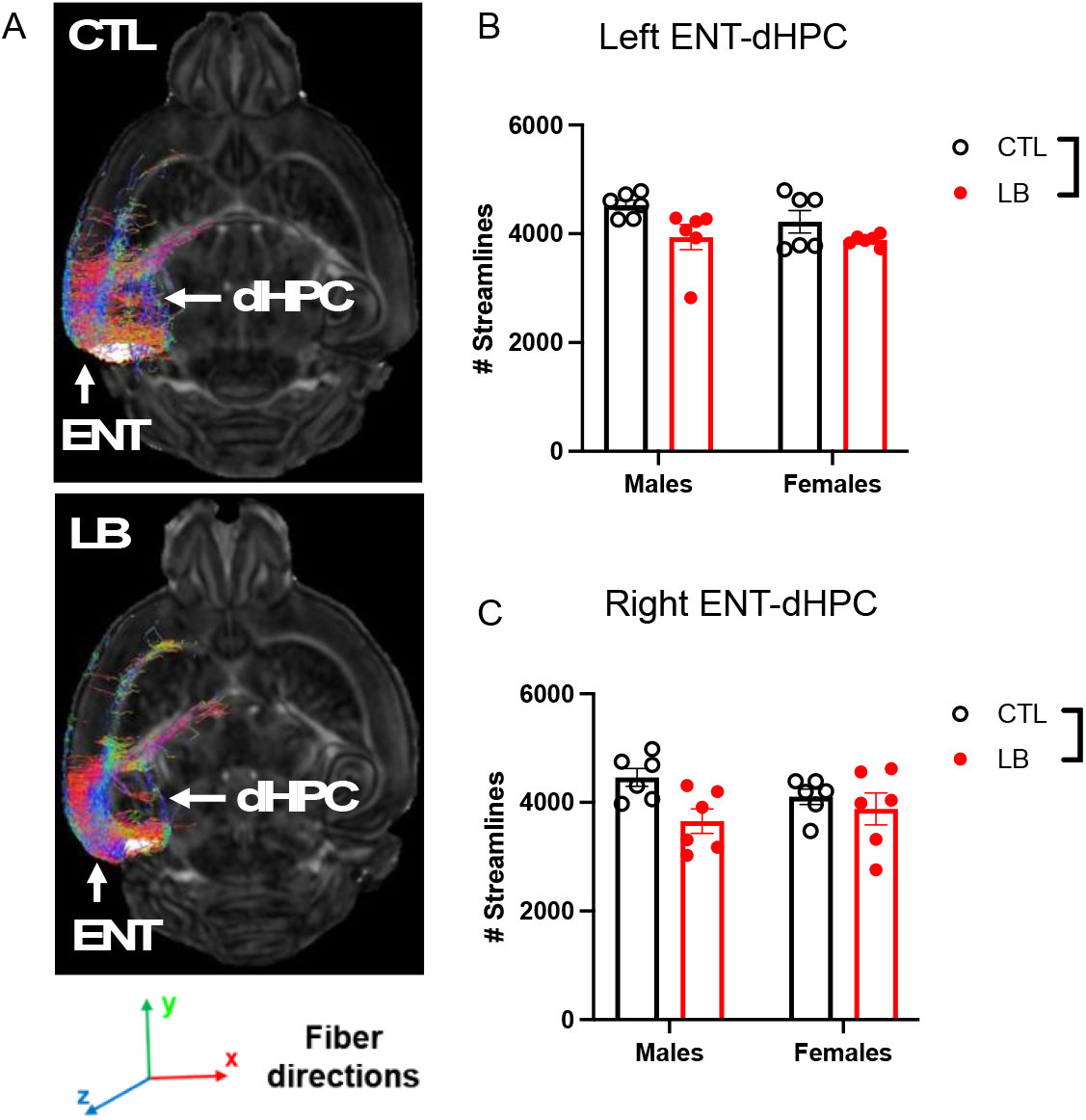
LB Reduces Structural Connectivity between the Entorhinal Cortex (ENT) and the Dorsal Hippocampus (dHPC) in P29 mice. Representative images (A) and quantification of ENT-dHPC tractography in the left (B) and the right (C) hemispheres. N = 6 mice per rearing and sex condition. Error bars represent mean ± SEM. *p < 0.05, **p < 0.01.

**Figure 8.**
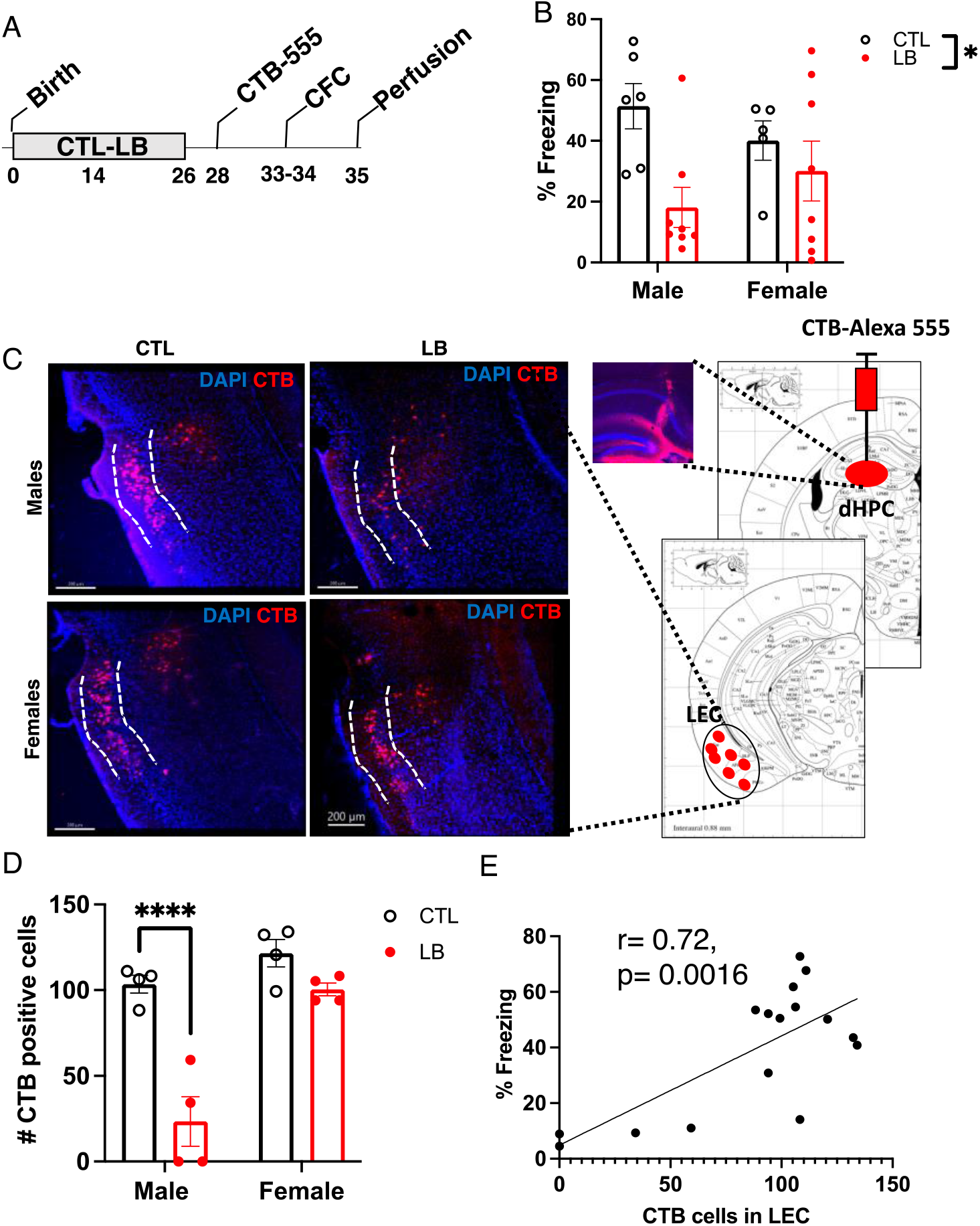
LB Causes Sex-Specific Deficits in Lateral Perforant Pathway Connectivity in Adolescent Mice. (A) CTL and LB mice were injected with the retrograde tracer Alexa 555-CTB into the left SLM at P28, tested in contextual fear conditioning (P33-34), and perfused to assess number of CTB positive cells in the LEC. (B) Contextual fear conditioning. (C) Schematic depiction of CTB injection at P28 and CTB labeling in the LEC at P35. (D) Number of CTB-positive cells in the LEC at P35. (E) Number of CTB-positive cells in the LEC correlates with contextual fear conditioning.

No significant effects of rearing, sex or interaction were seen for CTB labeling at the injection site (**Fig S5**). However, there was a significant interaction between rearing and sex for the number of CTB-positive cells in the LEC (F (1, 12) = 11.08, P = 0.0060). This interaction was due to a four-fold reduction in the number of CTB-positive cells in LB males (P < 0.0001) that was not observed in LB females (P = 0.22, **Fig 8C-D**). Moreover, the number of CTB-positive cells in the LEC was highly correlated with freezing behavior (**Fig 8E**). Together, these findings indicate that LB reduces connectivity between the entorhinal cortex and the dorsal hippocampus in a manner that correlates with deficits in contextual memory and is more pronounced in adolescent males.

Previous work has shown that reelin-positive cells project from the LEC to the dorsal hippocampus and that these projections are essential for contextual memory (23, 28, 51). To determine whether the CTB-positive cells in the LEC were reelin-positive, we repeated the CTB retrograde labeling using a second cohort of adolescent mice and replicated the sex-specific reduction in the number of CTB-positive cells in the LEC (Interaction: F (1, 13) = 10.76, P = 0.0060, **Fig 9A-B**). We confirmed that all CTB-positive cells were also reelin-positive, regardless of rearing and sex (**Fig 9A & C**), and found that the total number of reelin-positive cells in the LEC was reduced in LB mice, an effect that was more pronounced in males (rearing: F (1, 13) = 31.39, P < 0.0001, sex: (F (1, 13) = 25.50, P = 0.0002, interaction: (F (1, 13) = 3.399, P = 0.088, **Fig 9A & D**). The percentage of reelin-positive and CTB-negative cells was significantly higher in LB males compared to all other groups (interaction: F (1, 13) = 6.375, P = 0.025, **Fig 9 A & E**), indicating that most reelin-positive cells in the LEC fail to project to the dorsal hippocampus in LB males.

**Figure 9.**
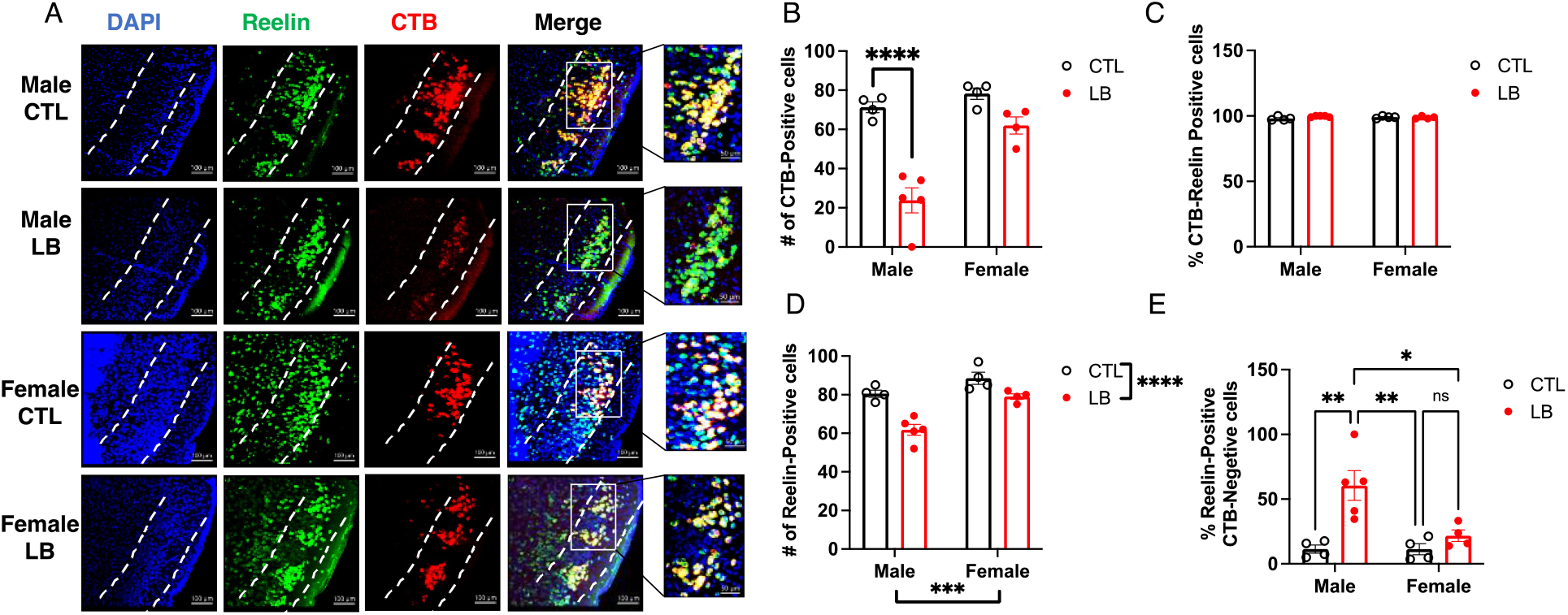
LB Reduces the Number of Reelin-Positive Projections in Males. (A) Representative images of Reelin and CTB positive cells in the LEC (Scale bars in Fig 9A are 50 microns). Higher magnification of rectangular areas of merged images are shown on the right. (B) The number of CTB-positive cells is reduced in the LEC of male LB mice. (C) All CTB-positive cells are reelin-positive regardless of rearing and sex condition. (D) LB reduces the number of reelin-positive cells in the LEC. (E). Effects of rearing and sex on the percentage of reelin-positive-CTB-negative cells in the LEC.

## DISCUSSION

This work provides several novel findings regarding the effects of LB, a commonly used mouse-model of ELA, on neurodevelopment and cognitive function in adolescent mice. First, we show that LB impairs connectivity between the LEC and the dorsal hippocampus in males and to a lesser extent in female mice, a finding that may explain why contextual freezing deficits are more severe in males. Second, adolescent LB mice showed hyperactivity and extensive cortical atrophy consistent with findings reported in children and adolescents exposed to severe early deprivation. Third, LB inhibited OPC differentiation in P17 pups and caused persistent hypomyelination in the SLM of adolescent mice, highlighting another layer of complexity by which early deprivation alters hippocampal function.

### LB Adolescent mice show Sex-Specific Deficits in Perforant Pathway Connectivity and Contextual Fear Conditioning

Exposure to traumatic brain injury in childhood disrupts entorhinal cortex-hippocamps connectivity and impairs episodic memory (29). Individuals exposed to childhood neglect/deprivation also show deficits in hippocampal dependent memory (33, 34) but a possible link to abnormal perforant pathway connectivity has not been reported. Here we show that the connectivity between the entorhinal cortex and the dorsal hippocampus is disrupted in a mice raised under impoverished conditions. Reduced connectivity was seen using dMRI tractography, retrograde tracing, and axonal staining in the SLM and was highly correlated with outcomes in the contextual fear conditioning.

Our data indicate that abnormal perforant pathway connectivity contributes to the more severe cognitive deficits found in adolescent LB male mice. This assertion is supported by work showing that these projections are essential for contextual memory (23, 28, 29, 51, 52) and the significant positive correlation between the number of these projections and contextual freezing. Reduction in reelin-positive cells in the LEC is more pronounced in LB males and is consistent with the volumetric changes seen with dMRI. However, the decrease in the number of reelin-positive cells is relatively small (roughly 25% reduction in males, Fig 9D), and it is the large percentage of reelin-positive/CTB-negative cells in the LEC of male LB mice that accounts for the striking sex differences in the number of CTB-positive cells. This is significant because it suggests that abnormal axonal pathfinding or axonal degeneration/retraction (53-55) may account for the inability of reelin-positive cells to maintain functional connections with the dorsal hippocampus in LB male mice. Future studies will determine whether the reduction in perforant pathway connectivity contributes to sex-specific changes in neurogenesis, synaptic pruning, and myelination, and whether strategies that augment these connections can normalize contextual memories in LB male mice.

### LB Replicates Key Features of Childhood Deprivation in Humans

Childhood neglect/deprivation is the most common form of early adversity (56, 57) leading to structural, behavioral, and cognitive deficits that are not typically seen in other forms of ELA such as physical and sexual abuse (30-32). Some of the key features of childhood neglect and deprivation include hyperactivity and cortical thinning (30-34, 49, 50), outcomes that have not been described yet in rodent models of ELA. The extended exposure to impoverished upbringing used in our LB paradigm coupled with the hyperactivity and extensive cortical atrophy found in adolescent LB mice, suggest that LB is a mouse model of early deprivation. One possible mechanism to explain the reduction in grey matter involves enhanced apoptosis in the cortex of LB mice. This process peaks between P2-P5, eliminates roughly 13% of all pyramidal cells, and is primarily driven by neuronal activity (58, 59). Impoverished conditions such as LB, coupled with erratic and fragmented maternal care (18), may decrease cortical activation during the first week of life, leading to enhanced cell death and reduced cortical volume. Clarifying the mechanisms responsible for the reduction in cortical volume and its impact on hyperactivity and cognition are important questions for future studies.

### LB Impairs Myelination in the SLM

Abnormal myelination is one of the most consistent findings in individuals exposed to early deprivation with similar findings reported in non-human primates and rodents (60). Only a few studies have examined the effects of LB on myelin development (35, 61) and the mechanisms responsible for these changes are yet to be clarified. Here we show that LB inhibits OPC differentiation and myelination in the SLM of P17 pups. This blockade is partly due to reduced axonal innervation, but the presence of hypomyelinated axons in the SLM suggests that other factors also contributed to reduced myelination. Since myelination is driven by neuronal activation (60), it is possible that the impoverished conditions associated with LB causes abnormally low levels of neuronal activation in perforant pathway terminals that further impair OPC differentiation. In addition, recent work has shown that LB reduces the expression of sodium channels in OPC located in the developing hippocampus (61). These channels may play a role in promoting OPC-differentiation in response to neuronal activity (62-64). Changes in microglial function may also contribute to blockade in OPC differentiation. For example, LB reduces expression of the receptor TREM2 in microglia (10), and low TREM2 levels are associated with abnormal OPC differentiation in a Cuprizone model of remyelination (65). LB also increases TNFα levels in microglia, which has been shown to inhibit OPC differentiation (66, 67).

The numbers of OPC and mature oligodendrocytes in the SLM were similar in adolescent CTL and LB mice, indicating that the blockade in OPC differentiation is transient. Nevertheless, levels of MBP staining in the SLM continued to be lower in adolescent LB mice suggesting that deficits in myelination persist at this age. Whether hypomyelination reflects a reduction in axonal availability or other mechanisms are questions that remain unresolved. Additional studies are needed to clarify whether agents that augment myelination (60) could recover myelination in the SLM and improve contextual fear learning in LB mice.

## CONCLUSIONS

Extended exposure to LB causes behavioral, cognitive, and structural changes that resemble those reported in children and adolescents exposed to early deprivation. These include hyperactivity, deficits in semantic memory, cortical atrophy, and abnormal myelination. A failure of reelin-positive cells to maintain functional connections with the dorsal hippocampus is likely to contribute to the more pronounced contextual-memory and myelination deficits found in adolescent LB male mice.

## ACKNOWLEDGEMENTS AND DISCLOSURES

We want to thank Dr. MacDowell Kaswan for helpful comments on the manuscript and the Yale Center for Genomic Analysis for the RNA sequencing. This work was supported by: NIMH R01MH119164, NIMH R01MH118332, and the Clinical Neuroscience Division of the VA National Center for PTSD. The authors declare no conflict of interest.

---

**Fig S1.**
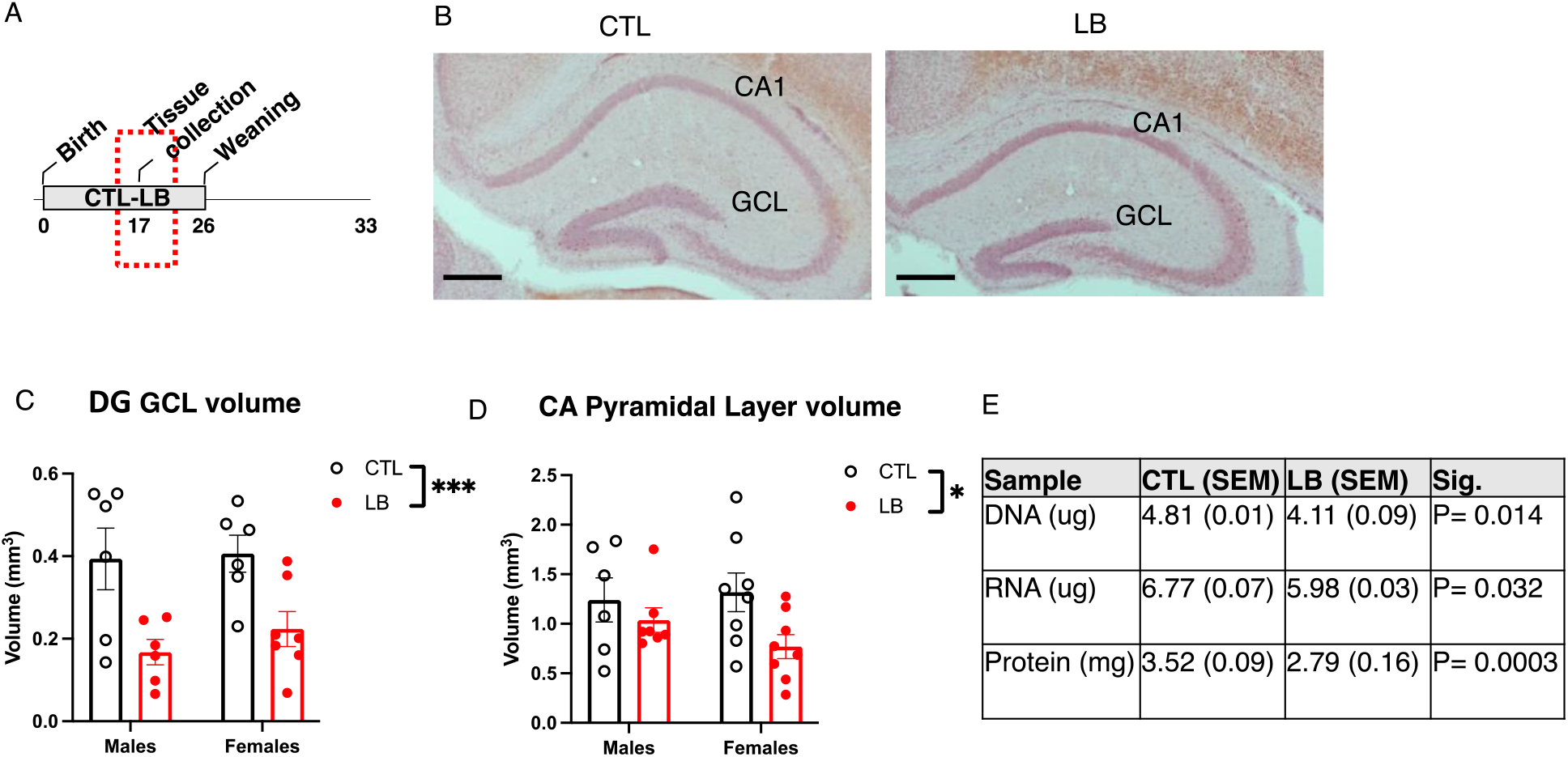
LB Impairs Hippocampal Development. (A) Litters were randomized to control (CTL) or limited bedding (LB) conditions at P0 and tissue was collected at P17. (B) Representative images of the dorsal hippocampus of P17 CTL and LB pups. LB reduces dentate gyrus (C) and Cornu Ammonis (D) in both male and female P17 pups. (E) DNA, RNA, and protein contents in the P17 hippocampus of CTL and LB pups. N = 6 mice per rearing and sex conditions. Scale bars in Fig B are 500um. Error bars represent mean ± SEM. *p< 0.05, ***p< 0.001.

**Fig S2.**
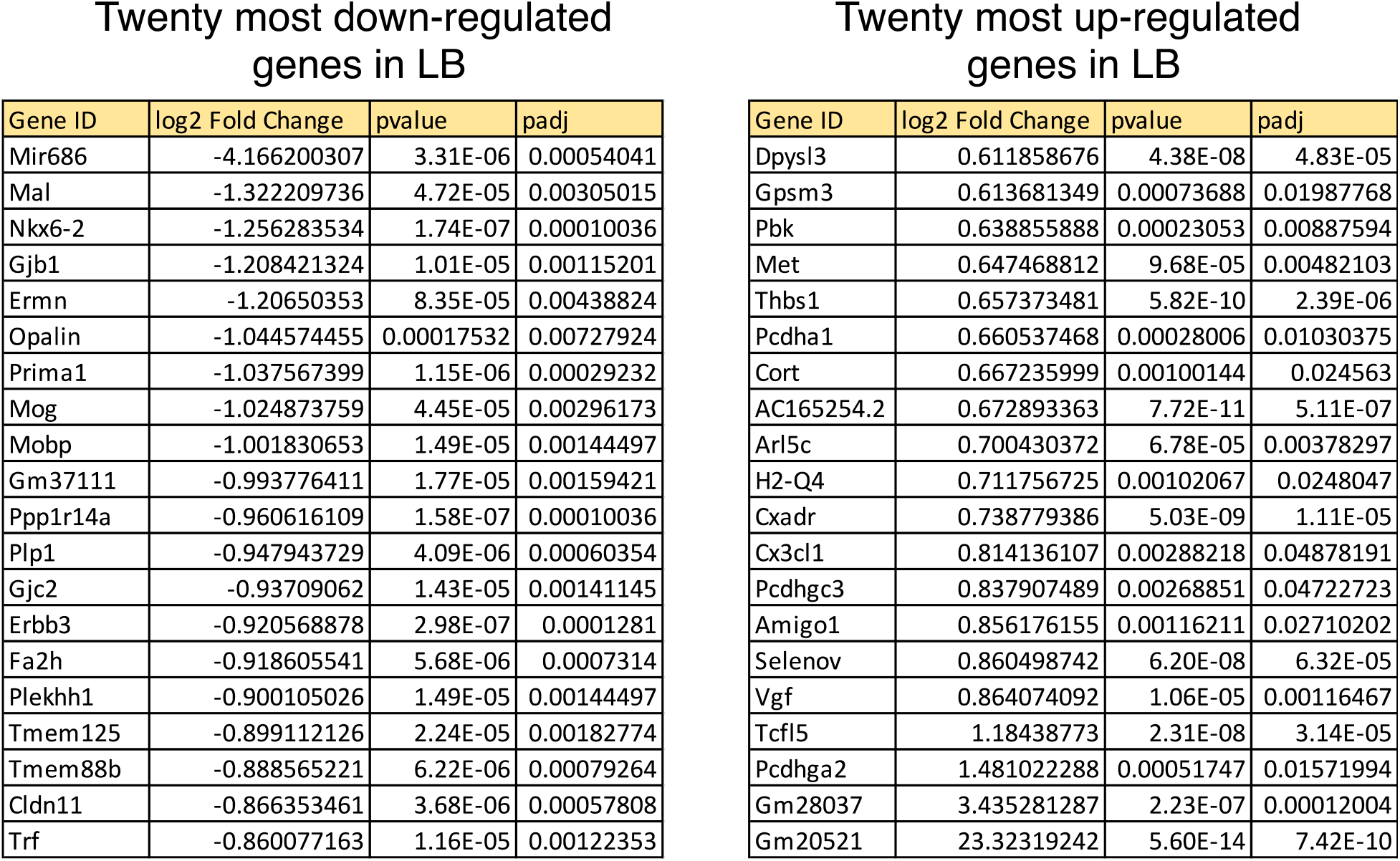
The Top 20 Most Downregulated and Upregulated genes in the P17 hippocampus. Padj is the adjusted p value after Benjamini-Hochberg correction for multiple comparisons.

**Fig S3.**
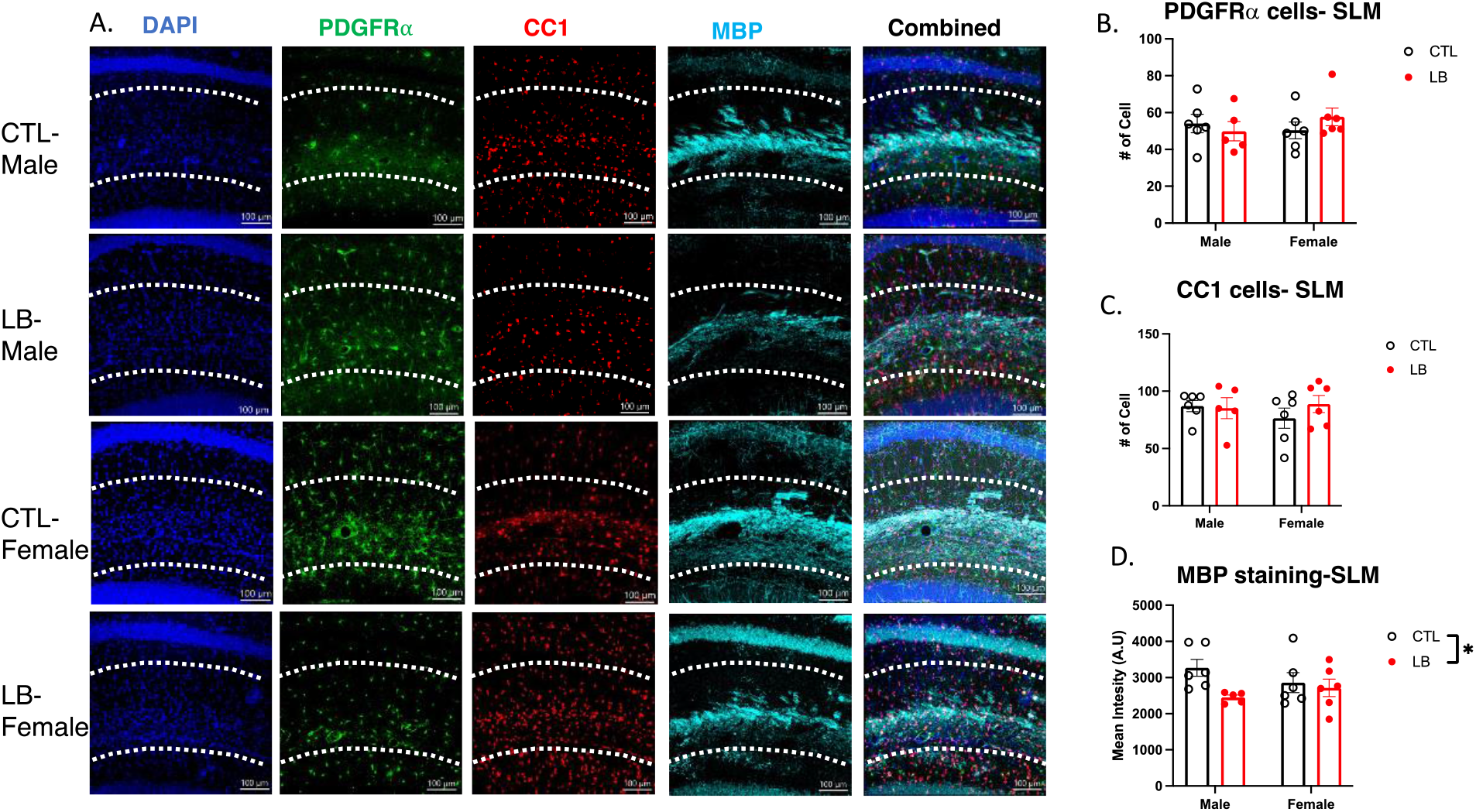
LB Impairs Myelination in the SLM of adolescent (P34) mice. (A) Representative confocal images of PDGFRα-positive OPC, CC1-positive mature oligodendrocytes, and MBP in the SLM of adolescent mice. Quantification of number of PDGFRα-positive OPC (B), CC1 mature oligodendrocytes (C) and MBP staining (D) in the SLM. Scale bars in Fig A are 100 microns. N = 5-6 mice per rearing and sex group. Error bars represent mean ± SEM. *p< 0.05.

**Fig S4.**
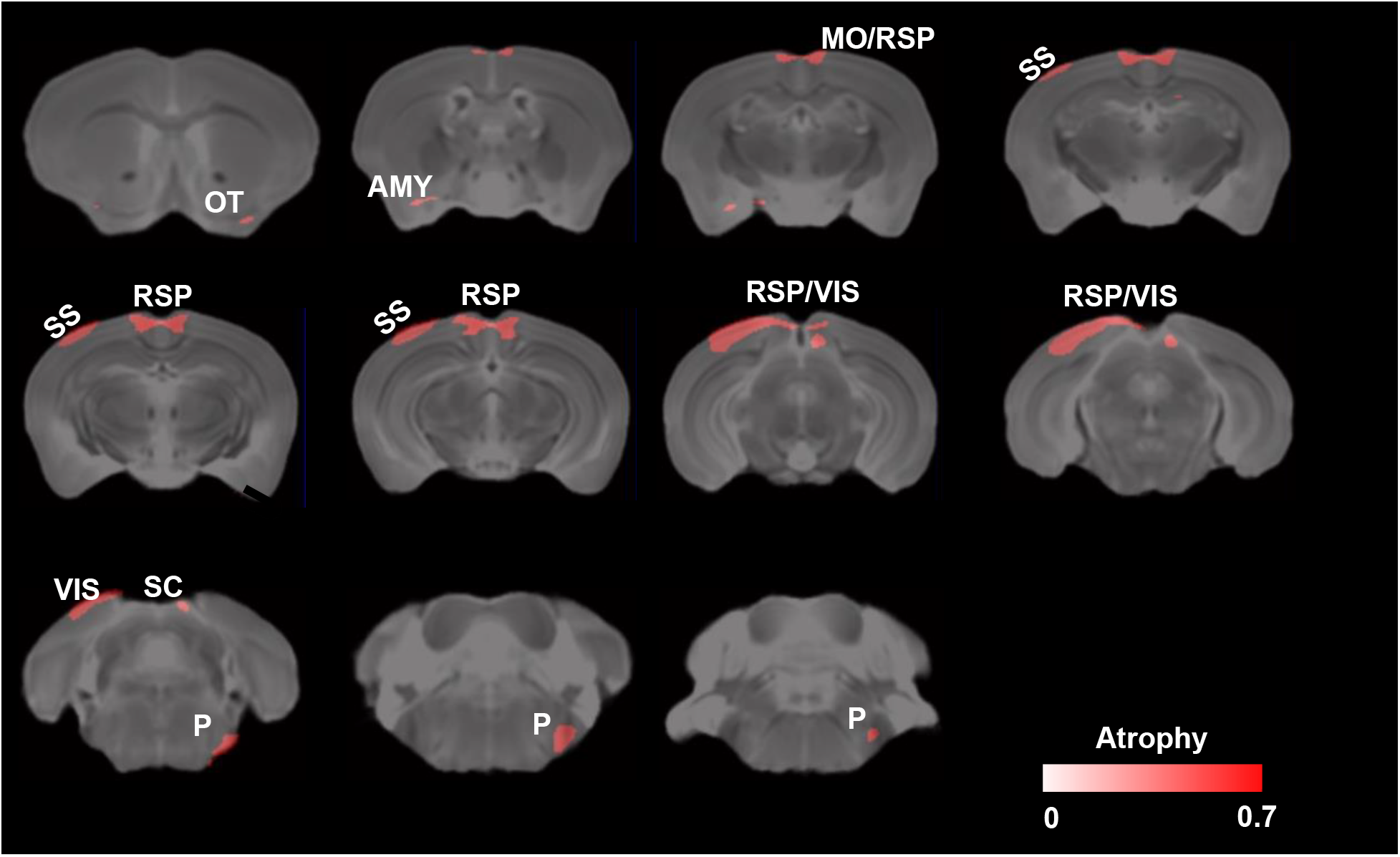
Main effect of sex on local volumetric changes in P29 adolescent mice. Minimal cluster size > 25 voxels, FDR< 0.1, p< 0.005. Areas with significant volume reduction in females are shown in red. Abbreviations: Amy: Amygdala, MO: Motor cortex, OT: Olfactory tubercle, P: Pons, RSP: Retrosplenial cortex, SC: Superior colliculus, SS: Somatosensory cortex, VIS: Visual cortex. N= 6 per rearing and sex group.

**Fig S5.**
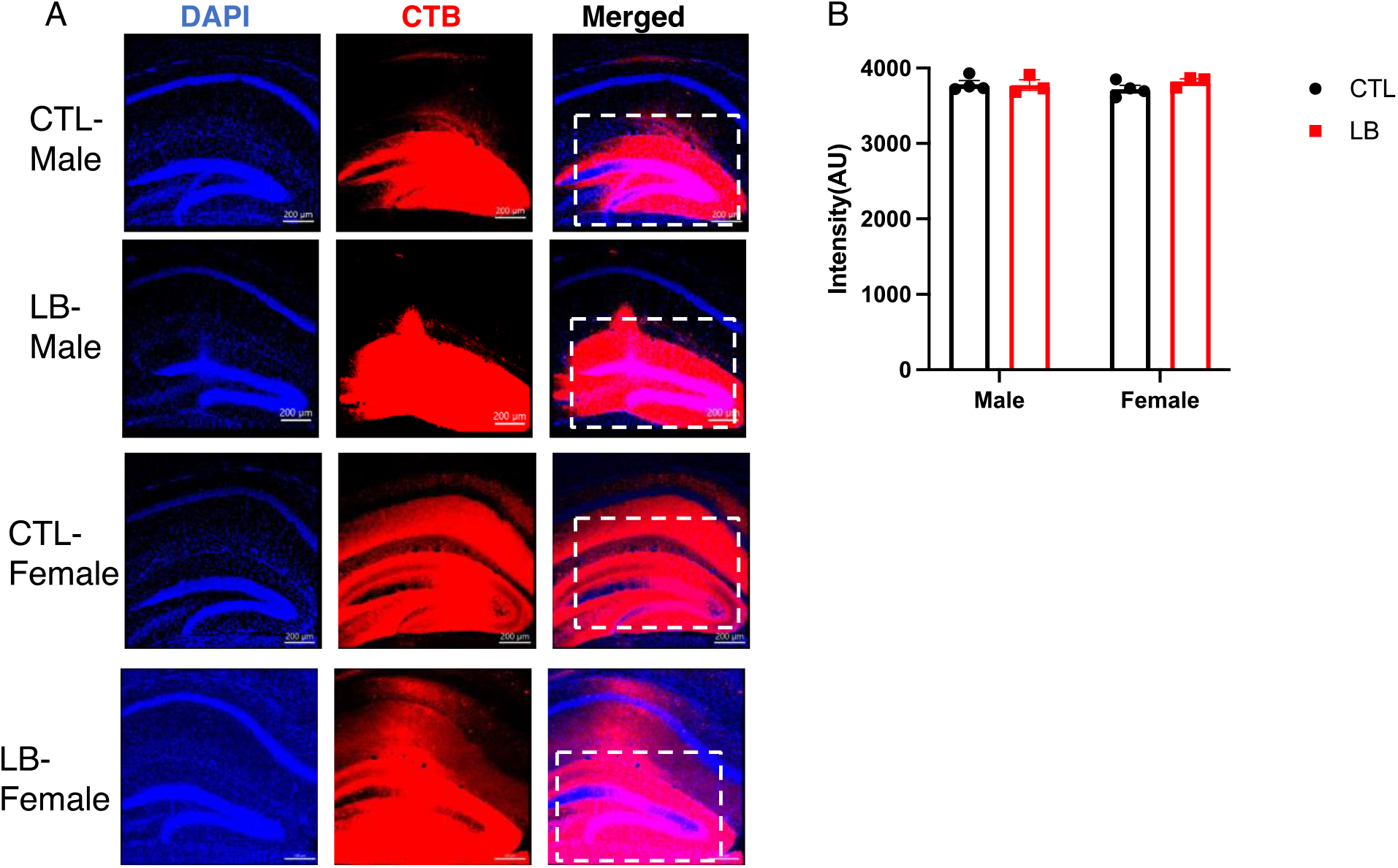
CTB Targeting of the Dorsal Hippocampus. (A) Representative images of CTB staining in the dorsal hippocampus of mice used to assess for retrograde labeling in Fig 9A. (B) Quantification of the signal found no significant effect of rearing F (1, 10) = 0.67, P= 0.43, sex F (1, 10) = 0.03, P= 0.86, or interaction F (1, 10) = 1.073, P= 0.32.

